# Calculation of sequence space coverage in a mutagenesis library

**DOI:** 10.64898/2026.06.15.729752

**Authors:** Alberto Florez Prada, Guido Uguzzoni, Darren J. Hart

## Abstract

Directed evolution requires screening of large mutagenesis libraries, but accurate calculation of library sizes needed to discover functional variants remains challenging. Existing models provide baseline estimates, yet current computational approaches for finding the best variants scale poorly with library complexity. Here, we introduce a scalable algorithmic framework to compute exact discovery probabilities in saturation mutagenesis libraries with no requirement for explicit sequence enumeration. By aggregating variants into a composition log–sum distribution and applying log-space convolution across randomisation blocks, it is possible to extend this to massive sequence spaces and mixed codon schemes.

By inverting these calculations, absolute mathematical ceilings for experimental design are established. Ultimately, this framework provides a rapid, quantitative tool to balance the statistical coverage-diversity trade-off within the limitations of laboratory screening.

Finally, this is implemented as an open-source web application (SSCC) that allows researchers to construct heterogeneous library designs and compute required sampling depths, coverage probabilities, and absolute randomisation limits.

## 1 Introduction

Directed evolution relies on the generation of molecular diversity through random or semi-rational mutagenesis, followed by screening for improved function. Saturation mutagenesis in which specific amino-acid positions are fully or partially randomised, offers precise control over diversity while maintaining manageable experimental quantities ^1^. Designing these libraries requires balancing coverage of the variant space with experimental feasibility: too small a library risks missing beneficial variants, while too large a library becomes unfeasible to construct or screen.

Early works established the foundational combinatorial and probabilistic formulations for estimating library degeneracy, completeness, and diversity, introducing Poisson approximations and analytical expressions for coverage probabilities ^1;2^. These models provided the first practical estimates of how many transformants are required to achieve complete coverage in random or cassette mutagenesis experiments. The conceptual distinction between degeneracy (the number of distinct sequences present in a library) and *diversity* (how much, on average, two randomly selected library members differ in sequence) clarified how redundancy in codon usage limits the effective search space of randomised libraries ^2^.

Building on these principles, subsequent tools such as GLUE-IT and PEDEL-AA extended coverage estimation to protein-level libraries by using algorithms to aggregate the probabilities of codon and amino-acid substitutions ^3^. These dynamic programming approaches enabled the calculation of expected library completeness and sampling distributions without enumerating all possible variants, a crucial advance for multi-site randomisation schemes.

In parallel, optimisation of codon randomisation schemes is a key experimental concept to improve coverage efficiency. Classical degenerate codons (NNN, NNK/NNS, NNB) produce highly biased amino-acid distributions and introduce unwanted stop codons. Other schemes, such as DKS, which has 12 codons and codes for 10 amino acids, do not introduce stop codons and they are highly biased towards hydrophobic residues. Kille et al. (2013) and others showed that rationally designed balanced codon sets, such as MAX or 20/20 libraries, substantially reduce redundancy and lower the number of clones required to achieve equivalent coverage ^4^. These efforts highlighted how the probabilistic structure of the codon distribution directly determines the attainable diversity and that analytical modeling is indispensable for optimising library construction.

Despite these improvements, most coverage models remain limited to expectations of completeness or relied on approximate occupancy formulas. Nov introduced the probability *T* (*k*) that at least one of the *k* top variants is present in a library of size *S* ^5^. It acknowledges that full coverage is rarely necessary in practice and that focusing on the most functional subset of variants drastically reduces library sizes required to achieve the desired phenotypic enhancements. However, exact computation still scaled poorly with library complexity and heterogeneous probability distributions, requiring heavy computational power beyond 4–5 positions.

Subsequent work on related questions confirmed the value of probabilistic modeling over brute-force enumeration. Sieber et al. (2015) presented a biomathematical description of peptide and display libraries based on expected coverage and relative efficiency metrics, showing that analytic expressions could accurately predict diversity across complex encoding schemes ^6^. In chemical and peptide libraries, MacConnell and Paegel (2017) similarly applied Poisson occupancy statistics to model sampling behavior and false discovery rates in massive combinatorial screens ^7^. Together, these studies established a consistent probabilistic language for describing library quality. However, to our knowledge, practical algorithms capable of handling arbitrary single or mixed codon schemes at large scale remain unavailable.

Despite the availability of theoretical models, a practical accessibility gap remains. While analytical expressions exist, researchers without a computational background lack user-friendly software to apply these models to complex, mixed-codon designs, often leaving them reliant on simplified rules of thumb or outdated estimation methods.

Here, we introduce a scalable algorithmic framework for computing expected top-*k* discovery probabilities (*T*_*k*_(*S*)) in saturation mutagenesis libraries without the need for explicit sequence enumeration. The method aggregates variants with identical probabilities into a compressed log-space composition distribution and combines heterogeneous randomisation blocks through mathematical convolution. This compositional–convolution approach perfectly reproduces the canonical analytical results of Nov (2012), while effortlessly scaling to billions of potential variants across arbitrary combinations of codon schemes. Additionally, we derive an exact analytical solution for libraries lacking codon bias (e.g., MAX). By inverting these coverage calculations, we establish practical mathematical ceilings for experimental design. Ultimately, this integration of exact analytical modelling with highly efficient computation delivers a unified, quantitative tool that enables researchers to design high-efficiency, low-bias mutagenesis libraries using only modest computational resources. These resources can be easily accesible through a standalone web implementation.

## 2 Materials and Methods

All computations were performed in Julia (version 1.10) using the packages `Roots` for root-finding, `SpecialFunctions` for log-factorial evaluation, and `CairoMakie` for figure generation. The analysis is distributed as a Jupyter notebook.

The computational pipeline proceeds in three stages. First, the function `compositions_ logdist` builds the base composition distribution for a small block of *m* ≤ 3 positions via a depth-first recursion in log-space, aggregating all compositions that share the same log-probability into a single entry and accumulating their log-multiplicities. Second, the convolution operator `convolve_logdists` and the exponentiation-by-squaring function `dist_power` combine base blocks to construct the full composition distribution for *L* randomised positions in O(log *L*) convolution steps, avoiding explicit enumeration of the exponential sequence space. Third, the functions `T1_from_dist` and `coverage_lower_bound` evaluate the discovery probability *T*_1_(*S*) and the full-coverage lower bound for a given library size *S* using exact binomial probabilities computed through the numerically stable identity (1 −*p*)^*S*^ = exp(*S* ln(1 −*p*)).

To construct Supplementary Table S1, the required library size *S* achieving a target probability was obtained by bisection root-finding (`Roots.Bisection()`) over the interval log_10_ *S* ∈ [0, 20]. Values exceeding 10^20^ are reported as *>* 10^20^.

### 2.1 Web implementation

To make these computations accessible, the algorithmic logic was ported into a client-side web application. The Sequence Space Coverage Calculator (SSCC) is built using standard web technologies (HTML, CSS, JavaScript) to ensure zero-dependency, local execution directly within the user’s browser, without requiring server-side processing.

SSCC is publicly available at https://github.com/florez-alberto/SSCC.

## 3 Results

### 3.1 Compositional representation of the sequence space

For each set of randomised positions, the probability of generating any sequence results from the product of the amino acid probabilities at each site. Instead of enumerating every possible sequence explicitly, we represent only how many times each amino acid appears, regardless of order. As we want to compute the multiplicity and the coverage of the sequence space, we can treat each site independently. We define the composition space, comprising all integer vectors **l** = (*l*_1_, *l*_2_, …, *l*_*A*_) whose entries sum to *L*, the number of randomised positions, where *A* is the number of amino acids possible given the randomisation scheme. This transformation allows to map the sequence space that is of order *A*^*L*^ (a factor that depends on the codon scheme) to a space of size 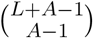.

Each composition represents all sequences that contain the same counts for each amino acid. The codon usage is then described by the probability of obtaining a specific sequence and its multiplicity. The codon scheme is defined by a probability vector **q** = (*q*_1_, …, *q*_*A*_) giving the probability of each amino acid. We defined the probability and the log-probability of a composition *l* as

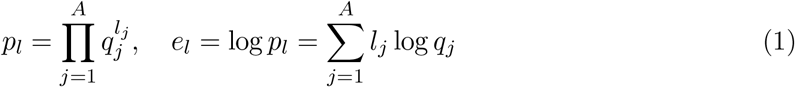

and the log-multiplicity of a composition *l* as

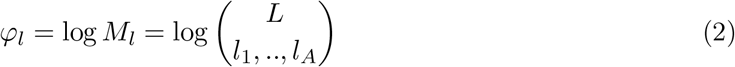

that represents the number of distinct sequences sharing a given composition, obtained via the multinomial coefficient.

The total number of variants is given by *n* = ∑_*l*_ *M*_*l*_. A composition space *D* is fully defined by the triple (*l, e*_*l*_, *φ*_*l*_).

### 3.2 Convolution of multiple codon schemes

The formulation above accounts for a specific mutagenesis scheme applied to *L* sites. When two different schemes are combined—*L*_1_ sites mutated with scheme **q**_**1**_ and *L*_2_ sites mutated with scheme **q**_**2**_—the probability and multiplicity of any sequence of length *L* = *L*_1_ + *L*_2_ in the composite scheme are obtained by multiplying the corresponding probabilities and multiplicities of the subsequences.

Calling *D* the composition space of the whole sequence and *D*_1_, *D*_2_ the composition space of the subsequences to merge we have *D* = *D*_1_ ⊕ *D*_2_ = (*l* = (*l*_1_, *l*_2_), *e*_*l*_ = *e*_*l*1_ + *e*_*l*2_, *φ*_*l*_ = *φ*_*l*1_ + *φ*_*l*2_).

### 3.3 Discovery probability: average probability to include a variant in the library

Given *D* and the total variant count *n*, the probability that a specific variant appears at least once in a library of size *S* is 1 − (1 −*p*)^*S*^. Averaging over all variants yields the expected discovery probability:

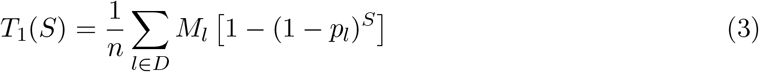

### 3.4 Codon schemes without codon bias (MAX)

In the case of a codon scheme without codon bias, all amino acids have the same probability *q*_*a*_ = 1*/A* and all variants have the same probability *p*.

In that case, we can analytically compute the discovery probability 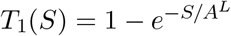.

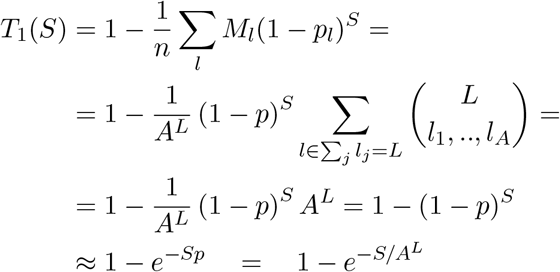

Where we used 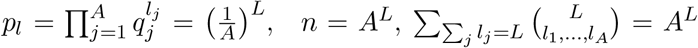, and (1 − *p*)^*S*^ ≈ *e* ^−*Sp*^ for *S* large.

### 3.5 General Codon schemes

#### Representation of codon schemes

A codon scheme can be characterized by how many amino acids share the same number of encoding codons. Let *C*_*i*_ denote the number of amino acids that are encoded by exactly codons, where *i* = 1,…, *i*_max_. These counts satisfy ∑_*i*_ *C*_*i*_ = *A* (the number of amino acids, typically 20) and ∑_*i*_ *i C*_*i*_ = *N*_*c*_ (the total number of sense codons).

Under uniform codon sampling, an amino acid encoded by *i* codons appears with probability *p*^(*i*)^ = *i/N*_*c*_.

#### Composition representation

A sequence of *L* randomized positions can be summarized by the vector 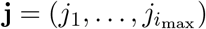, where *j*_*i*_ is the number of positions occupied by amino acids of the *i*-codon class. This vector satisfies ∑_*i*_ *j*_*i*_ = *L*.

The number of distinct sequences sharing the same composition **j** is:

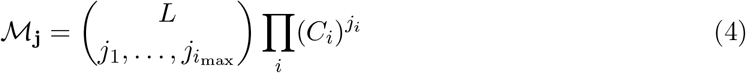

The multinomial coefficient counts the arrangements of codon classes across positions, while 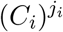 counts the choices of amino acids within each class (with repetition allowed, since the same amino acid can appear at multiple positions).

#### Discovery probability

The probability associated with composition **j** is 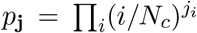. The expected coverage after sampling *S* library members is:

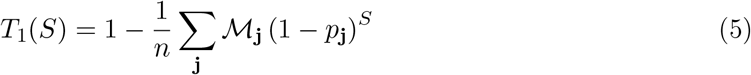

where *n* = *A*^*L*^ is the total number of amino acid sequences. For large *S*, using (1 − *p*)^*S*^ ≈ *e*^−*Sp*^:

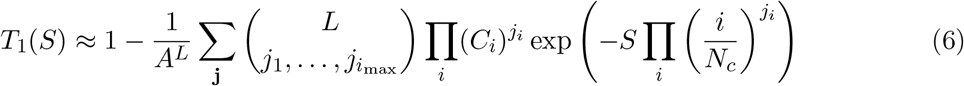

Unlike the simpler case of single-position coverage, this expression does not reduce to a closed form because the exponential of a product does not factorize. However, the computational advantage remains: the sum runs over composition vectors **j** satisfying ∑_*i*_ *j*_*i*_ = *L*, whose number grows as 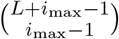 —combinatorially in *L* rather than exponentially as *A*^*L*^. This compression enables efficient numerical evaluation even for large libraries.

### 3.6 General codon schemes and log-space convolution

#### Base compositional representation

Instead of enumerating the full combinatorial space of *L* positions simultaneously, we define a scalable approach starting from a small block of *m* positions (e.g., *m* ≤ 3). A specific codon scheme (e.g., NNS, DKS) is characterized by a probability vector **q** = (*q*_1_, …, *q*_*A*_) defining the likelihood of obtaining each of the *A* amino acids.

For this base block of length *m*, we compute the composition space *D*_*base*_. Each unique composition within this block is compressed into a state defined by its log-probability 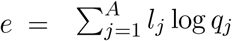 and its sequence multiplicity 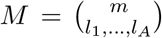. Identical log-probabilities resulting from degenerate codon frequencies are aggregated, naturally compressing the state space without requiring manual grouping of codon classes.

#### Exponentiation by squaring via convolution

To evaluate massive library sizes without facing a combinatorial explosion, we apply the convolution operator defined in Section 2.2. The convolution of two independent sequence blocks *D*_1_ (of length *L*_1_) and *D*_2_ (of length *L*_2_) yields a combined composition space *D*_*new*_ of length *L*_1_ + *L*_2_. For every pair of states *u* ∈ *D*_1_ and *v* ∈ *D*_2_, the merged state is computed in log-space:

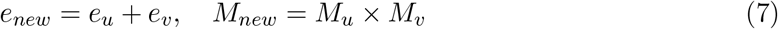

By utilizing exponentiation by squaring, a library of length *L* can be built in O(log *L*) convolution steps rather than scaling with the exponential sequence space O(*A*^*L*^) or the full multinomial composition space. For instance, computing a sequence of *L* = 12 positions requires only the base computation of *D*_3_, followed by two convolution squarings (*D*_6_ = *D*_3_ ⊕ *D*_3_, and *D*_12_ = *D*_6_ ⊕ *D*_6_).

#### Heterogeneous randomisation blocks

This convolutional framework natively supports heterogeneous library designs, where different *l* positions are subjected to different mutagenesis strategies. A library consisting of *L*_1_ positions mutated with scheme A and *L*_2_ positions mutated with scheme B is simply the convolution of their respective distributions:

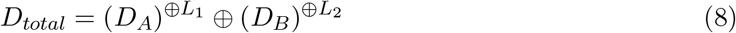

#### Exact binomial expected coverage

Once the final distribution *D*_total_ is computed, the expected coverage *T*_1_(*S*) for a sampled library of size *S* is evaluated. Rather than relying on a continuous Poisson approximation (*e*^−*Sp*^), which introduces numerical artifacts at extreme sampling depths due to the nature of computer calculations, we compute the exact discrete binomial probability of missing a variant, (1 −*p*)^*S*^. To maintain floating-point stability when *S* ≫ 10^8^ and *p*→ 0 (where 1 −*p* rounds exactly to 1.0 in standard 64-bit floating-point arithmetic), we utilize the mathematically equivalent but numerically stable transformation exp(*S* ln(1 −*p*)). In our computational implementation, the ln(1− *p*) term is evaluated using the standard log1p(-p) routine, which computes ln(1 + *x*) accurately even for vanishingly small values of *x*. This ensures the calculation:

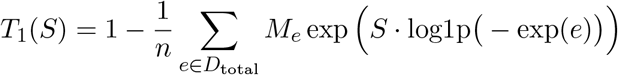

remains perfectly stable, allowing the framework to scale to sequence spaces and library sizes exceeding 10^20^ without catastrophic cancellation or underflow.

### 3.7 Top-*k* discovery probability

A natural extension of the coverage metric *T*_1_(*S*) is the probability of observing at least one among *k* target variants. However, computing this quantity exactly would require specifying the identities and individual sampling probabilities of the *k* target variants. In the absence of prior knowledge about which variants are functionally relevant, a reasonable approach is to average over all possible choices of *k* variants drawn uniformly at random from the sequence space. Under this interpretation, and assuming the *k* variants are chosen independently, the probability of missing all of them factorizes:

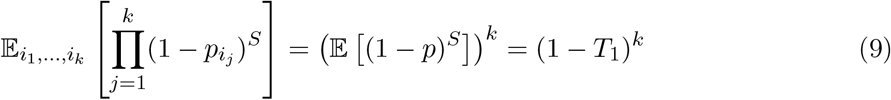

The expected discovery probability for *k* randomly chosen variants is therefore:

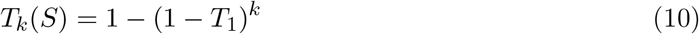

This relationship demonstrates how focusing on a subset of functional variants drastically shifts the required screening capacity. As *k* increases, high discovery probabilities can be achieved even when the physical sampling depth S is a fraction of the total sequence space *A*^*L*^(Supplementary Figure S2A).

By applying this framework, we calculated the exact library size requirements (*S*) to achieve 95% and 99% discovery probabilities across a wide range of *k* and *L* values (Supplementary Table S1). The compositional–convolution algorithm perfectly reproduces previously published exact analytical values for small-scale randomisations (*L* ≤ 3)^5^, while effortlessly extending the computation to massive sequence spaces (*L* ≥ 10) and mixed codon schemes that were previously computationally inaccessible.

### 3.8 Max *L* discovery probability

To establish practical limits for library design, we invert the coverage calculation: given a physical screening capacity *S* and a minimum acceptable discovery probability *T*_*target*_ what is the maximum number of positions L that can be randomised.

Because the accessible sequence space *A*^*L*^ grows exponentially with *L*, the expected coverage *T*_*k*_(*S*) strictly decreases as *L* increases for a fixed *S*. We determine this absolute boundary computationally by evaluating *T*_*k*_(*S*) across discrete, increasing integer values of L until the coverage drops below the *T* target.

This discrete step-function defines the absolute mathematical ceiling for an experiment. Relaxing the target discovery probability (*T*_target_) from a stringent 99% down to 50% yields surprisingly marginal gains; it typically expands the maximum permissible *L* by no more than a single position (e.g. in Supplementary Figure S2B). However, evaluating this boundary across different codon schemes reveals that optimised, stop-codon-free strategies (such as DKS and MAX) stretch this ceiling considerably (Supplementary Figure S2C). By drastically reducing combinatorial redundancy, these efficient schemes allow experimentalists to target significantly more positions simultaneously at the exact same sampling depth compared to standard NNN or NNK/NNS randomisation.

### 3.9 Practical library design with SSCC

To facilitate experimental design, we implemented the compositional-convolution algorithm in the SSCC web tool. The graphical interface allows users to dynamically calculate required library sizes (*S*), expected discovery probabilities (*T*_*k*_), and maximum randomisable positions (*L*). For instance, calculating the required *S* for an 8-position library using an NNS scheme versus a mixed NNS/DKS design can be performed instantaneously via the interface, explicitly illustrating the shifts in sequence space coverage. A representative view of the user interface is provided in **Supplementary Figure S3**.

## 4 Discussion

### 4.1 The coverage-diversity trade-off and selection power

A practical application of this framework is determining the maximum number of positions *L* that can be randomised while maintaining a target discovery probability for a given screening capacity *S*. However, this formulation obscures a fundamental trade-off: increasing *L* expands the accessible sequence space exponentially (*A*^*L*^), potentially including higher-fitness variants, but simultaneously reduces the coverage achievable at fixed *S*.

We emphasise that calculating the maximum *L* is not intended to serve as a direct comparison of biological fitness between different library sizes. Rather than predicting biological superiority, determining the maximum *L* serves as a strict boundary for experimental design: it should prevent researchers from designing libraries so large that undersampling guarantees the failure of the experiment. By performing an inverse calculation, we can establish the baseline, defining exactly how far a researcher can push the combinatorial space before the target discovery probability mathematically collapses. The experimentally relevant quantity depends on balancing this statistical ceiling against how fitness is distributed across the sequence space, as we discuss below.

Comparing libraries with different *L* is conceptually subtle. The sequence space at *L*_1_ *< L*_2_ is embedded in the larger space at *L*_2_, with the additional positions fixed at wild-type. A variant ranked first at *L*_1_ will have a different ranking in the *L*_2_ space, where it competes with sequences carrying additional mutations. The experimentally relevant quantity is the expected fitness of the best observed variant, which depends on both how many unique sequences are sampled and how fitness is distributed across the sequence space.

Under a simple additive model where each mutation contributes an effect drawn from a distribution with mean *µ <* 0 (most mutations are deleterious) and *σ*^2^ as the variance ^8^, the expected maximum fitness among *n*_eff_ observed sequences scales approximately as:

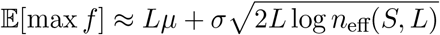

where 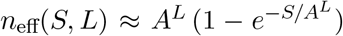 is the expected number of unique variants observed. This equation highlights two competing forces: a linear mutational load (*Lµ*) that depresses the mean fitness of the library, and a variance expansion 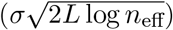 that stretches the fitness distribution into higher extremes. This expression exhibits a maximum at an intermediate *L*^∗^ that balances the deteriorating mean fitness against the expanding variance and sample size. However, the additive model ignores epistasis and requires parameters that are rarely known a priori.

To leverage the variance expansion both the physical evaluation (*S*_evaluation_) and the final screening or readout (*S*_screening_) capacities must be large, which is typically achieved by coupling display technologies with Next-Generation Sequencing (NGS) where *S*_screening_ ≈ *S*_evaluation_. At the same time, the design must respect the transformation efficiency (*S*_transformation_), which serves as the physical ceiling. For yeast display, *S*_transformation_ is typically 10^5^–10^6^, phage display 10^8^–10^9^, and ribosome display 10^12^–10^13^. In any robust design, *S*_transformation_ ≥ *S*_evaluation_.

### 4.2 Assumptions and biological realities

The top-*k* discovery metric assumes that the target functional variants are distributed uniformly and independently across the sequence space. In biological systems, high-fitness variants are often clustered due to epistasis and shared structural constraints. If a functional family of variants relies heavily on a codon composition that is rare within the chosen scheme, the true probability of capturing at least one of them may deviate from the theoretical uncorrelated estimate.

Second, the model establishes baselines using idealised probability vectors for degenerate codons (e.g., exactly 1/32 for each NNK codon). However, commercial oligonucleotide synthesis—particularly degenerate hand-mixing—is susceptible to coupling efficiency variations, meaning the empirical distribution is rarely perfectly uniform. A key advantage of the compositional–convolution algorithm presented here is that it is not restricted to idealised formulas. Because the framework accepts any arbitrary probability vector **q**, experimentalists can directly input empirical codon frequencies derived from NGS quality control data. This allows the tool to seamlessly model expected coverage under real-world synthesis biases.

### 4.3 Practical guidelines

In the absence of full fitness landscape information, experimentalists can consider two limiting criteria for library design. Fixing an absolute target *k* (e.g., *k* = 100) across all *L* is a conservative approach: it favours a smaller *L* with high coverage, ensuring that the best variants within a restricted sequence space are reliably found. Conversely, fixing a relative fraction *k/A*^*L*^ = *f* (e.g., targeting the top 0.001% of the landscape) favours a larger *L*: it searches for sequences that are highly exceptional relative to their own space, accepting much lower absolute coverage to do so. Neither criterion is universally correct; the choice dictates whether the biological objective requires multiple simultaneous mutations to achieve a functional gain.

As a practical guideline: when preliminary data (such as single-mutant fitness measurements) are available, estimating the mean mutational effect *µ* and variance *σ* provides a rational, data-driven basis for choosing *L*. When such data are lacking, a conservative approach favouring high coverage is generally preferable, as finding the best variant in a small, fully sampled space is more reliable than severely undersampling a vast one. Pushing for a larger *L* should be reserved for cases where structural or biological reasoning strongly suggests that complex epistatic combinations are strictly necessary for the desired phenotypic improvement.

### 4.4 Sequence Space Coverage Calculator (SSCC)

We implemented this algorithmic framework into an open-source, client-side web application called the Sequence Space Coverage Calculator (SSCC). The tool provides a graphical interface allowing researchers to calculate *S, T*_*k*_, and *L* dynamically. Users can model heterogeneous libraries by assembling modular sequence blocks of different lengths and predefined codon schemes (NNN, NNS/NNK, DKS, and MAX). By executing the log-space compositional convolution and exact binomial evaluations natively in JavaScript, the application ensures rapid, mathematically precise computations of massive sequence spaces directly within the browser, eliminating the need for local computational environments. Furthermore, because the underlying logic accepts arbitrary probability arrays, the tool’s modular architecture readily allows for future adaptations, such as incorporating empirical next-generation sequencing (NGS)-derived codon distribution frequencies or custom rationally designed codon mixtures.

## Supporting information

Jupyter notebook code

## 5 Code Availability

The Julia source code used for theoretical validation and figure generation, in Jupyter notebook, is publicly available at https://github.com/florez-alberto/SSCC. The JavaScript source code for the SSCC web application is also open-source and hosted within the same repository. The web tool can be accessed directly at https://florez-alberto.github.io/SSCC/.

## 6 Acknowledgements

AFP and DH received funding from the ANR FluPept project ANR-21-CE18-0024 and the France 2030 PUI program of Université Grenoble Alpes (UGA). IBS acknowledges integration into the Interdisciplinary Research Institute of Grenoble (IRIG, CEA). AFP received funding from the École doctorale Chimie et sciences du vivant (EDCSV) – UGA. GU gratefully acknowledges financial support from the IRIG/CEA FAST program (Fond d’Action Sciences et Techniques) for young researchers.

## 8 Supplementary Information

**Supplementary Table S1:**
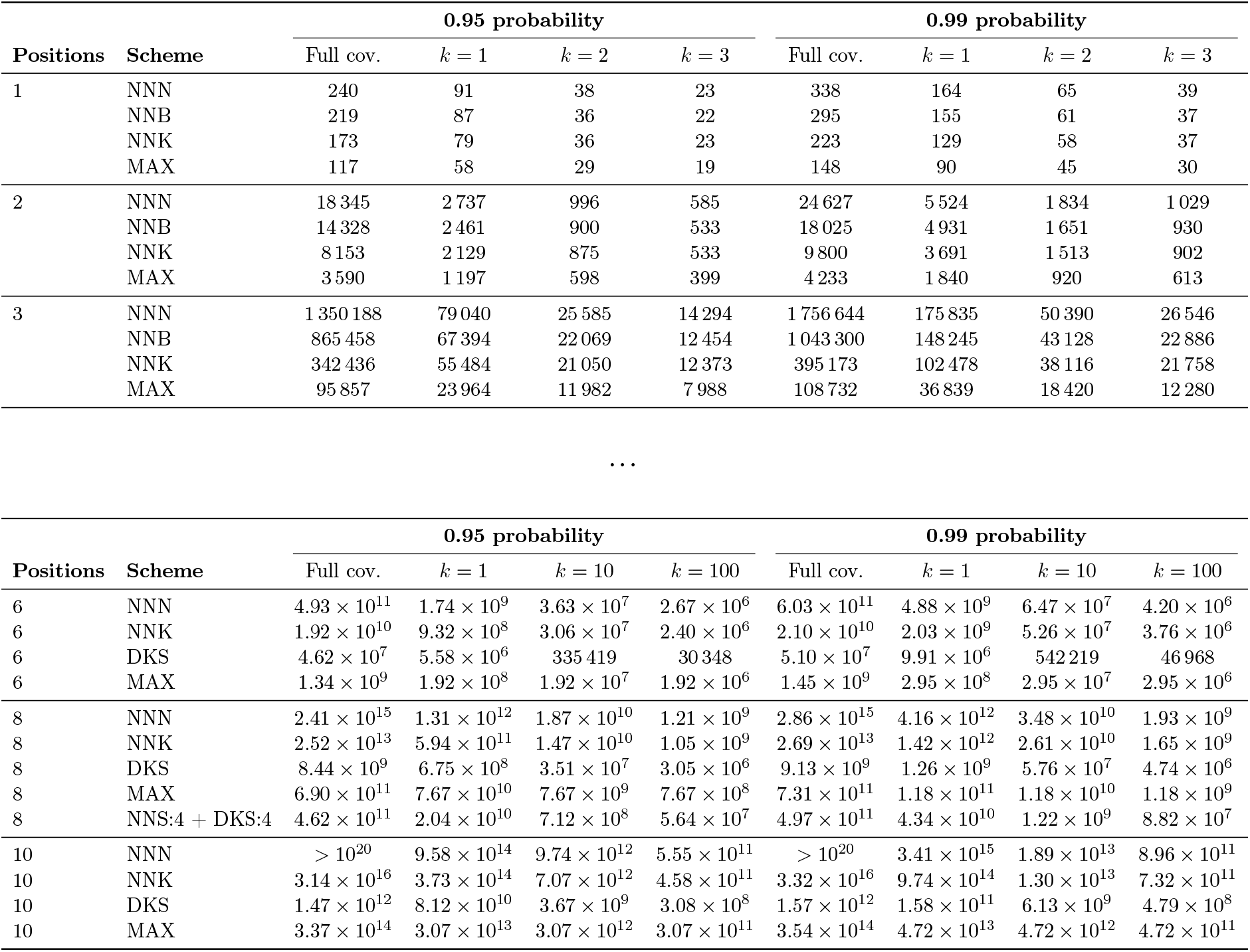
Required library sizes for full coverage and top-*k* discovery across various codon schemes. Calculated physical library sizes (*S*) necessary to achieve 0.95 and 0.99 discovery probabilities. The upper section (*L* = 1, 2, 3; target pool sizes *k* = 1, 2, 3) serves as a validation set, perfectly reproducing the exact analytical results previously established by Nov (2012). The lower section (*L* = 6, 8, 10; target pool sizes *k* = 1, 10, 100) demonstrates the scalability of the compositional–convolution algorithm, evaluating massive sequence spaces and heterogeneous randomisation blocks (e.g., the *L* = 8 mixed NNS/DKS scheme) that are otherwise computationally inaccessible via brute-force enumeration. Values requiring experimentally unfeasible library sizes exceeding 10^20^ transformants are indicated as *>* 10^20^.

| Positions | Scheme | 0.95 probability |  |  |  | 0.99 probability |  |  |  |
| --- | --- | --- | --- | --- | --- | --- | --- | --- | --- |
| | | Full cov. | $k = 1$ | $k = 2$ | $k = 3$ | Full cov. | $k = 1$ | $k = 2$ | $k = 3$ |
| 1 | NNN | 240 | 91 | 38 | 23 | 338 | 164 | 65 | 39 |
|  | NNB | 219 | 87 | 36 | 22 | 295 | 155 | 61 | 37 |
|  | NNK | 173 | 79 | 36 | 23 | 223 | 129 | 58 | 37 |
|  | MAX | 117 | 58 | 29 | 19 | 148 | 90 | 45 | 30 |
| 2 | NNN | 18 345 | 2 737 | 996 | 585 | 24 627 | 5 524 | 1 834 | 1 029 |
|  | NNB | 14 328 | 2 461 | 900 | 533 | 18 025 | 4 931 | 1 651 | 930 |
|  | NNK | 8 153 | 2 129 | 875 | 533 | 9 800 | 3 691 | 1 513 | 902 |
|  | MAX | 3 590 | 1 197 | 598 | 399 | 4 233 | 1 840 | 920 | 613 |
| 3 | NNN | 1 350 188 | 79 040 | 25 585 | 14 294 | 1 756 644 | 175 835 | 50 390 | 26 546 |
|  | NNB | 865 458 | 67 394 | 22 069 | 12 454 | 1 043 300 | 148 245 | 43 128 | 22 886 |
|  | NNK | 342 436 | 55 484 | 21 050 | 12 373 | 395 173 | 102 478 | 38 116 | 21 758 |
|  | MAX | 95 857 | 23 964 | 11 982 | 7 988 | 108 732 | 36 839 | 18 420 | 12 280 |

**Supplementary Figure S2.**
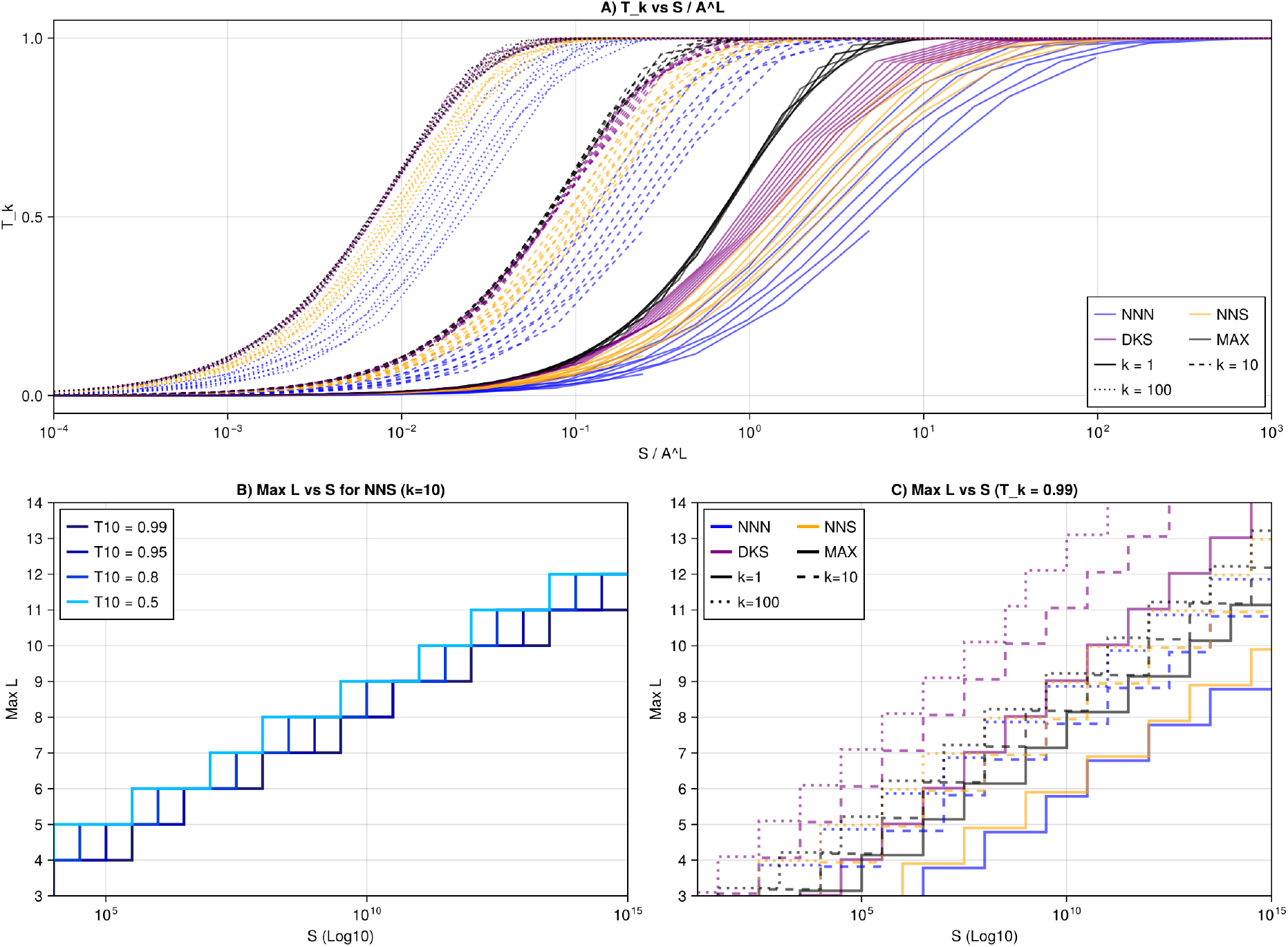
Discovery probabilities and absolute mathematical limits of library randomisation. **(A)** Expected top-*k* discovery probability (*T*_*k*_) plotted against the normalised screening capacity (*S/A*^*L*^). Curves illustrate the scaling behaviour of different codon schemes across sequence lengths *L* = 6, 8, 10 and fixed top *k* = 1, 10, 100 variants. **(B)** The maximum number of randomisable positions (Max *L*) using the NNS codon scheme to find at least one of *k* = 10 variants across varying target probability thresholds (*T*_10_). Relaxing the acceptable probability from a stringent 0.99 down to a 0.50 coin-flip yields only marginal gains in *L*. **(C)** The ceiling for experimental design (Max *L*) at a strict 0.99 discovery probability (*T*_*k*_ = 0.99), comparing classical (NNN, NNS) and optimised (DKS, MAX) codon schemes. Stop-codon-free, low-bias schemes drastically reduce combinatorial redundancy, raising the absolute ceiling for simultaneous mutations at any given sampling depth *S* (x-axes for B and C are shown on a log_10_ scale).

**Supplementary Figure S3.**
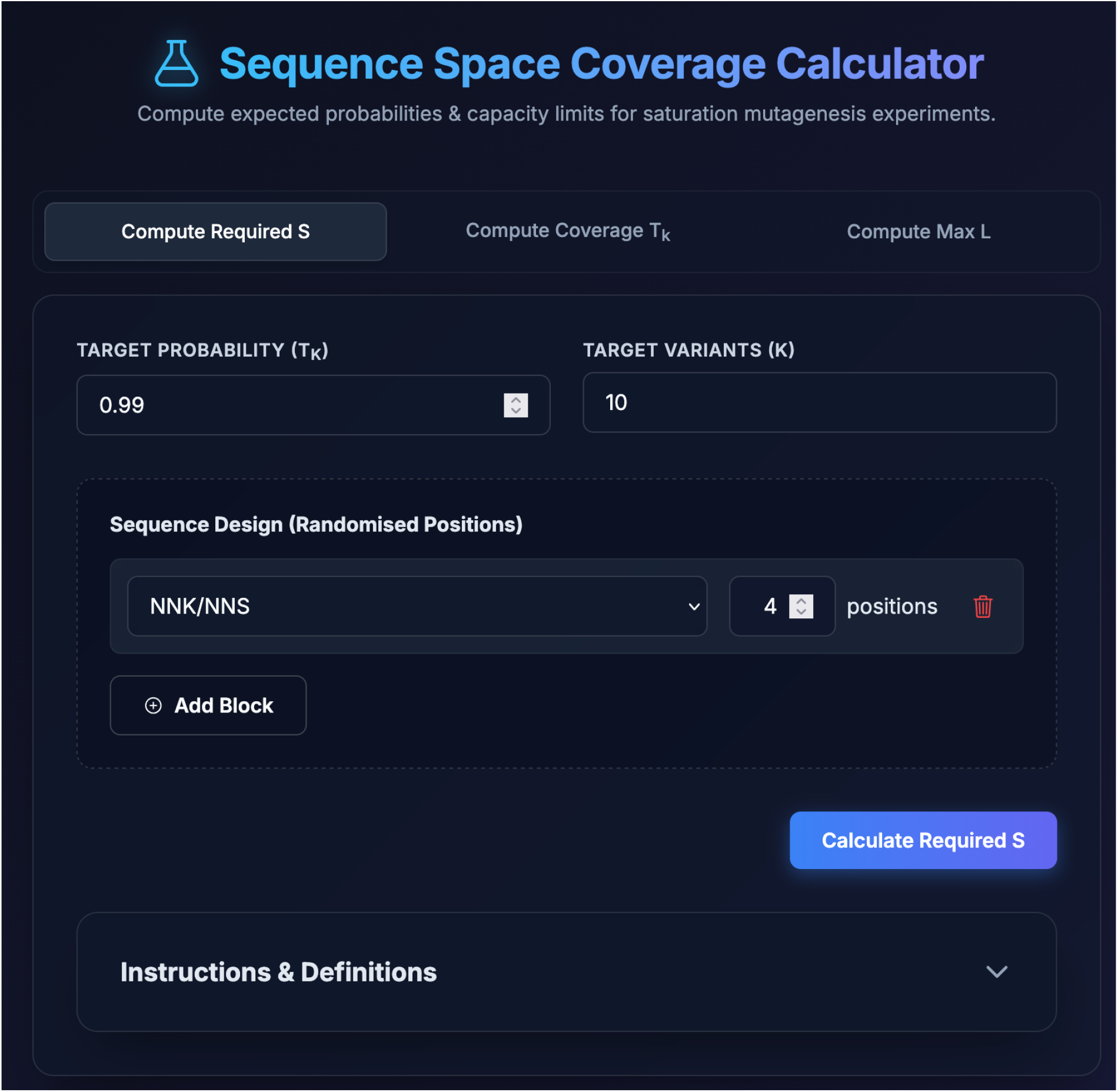
The Sequence Space Coverage Calculator (SSCC) web interface. The graphical user interface allows researchers to construct heterogeneous saturation mutagenesis libraries using modular sequence blocks. Users can specify target probabilities (*T*_*k*_) and the number of target variants (*k*) to dynamically compute the required experimental screening capacity (*S*), expected sequence space coverage, or the maximum number of randomisable positions (*L*). By executing calculations natively within the browser, the tool enables rapid experimental design without requiring local computational environments.

