## Supplementary material for "Calculation of sequence space coverage in a mutagenesis library": Jupyter notebook code

### Supplementary_code

June 10, 2026

[1]:
```
using Pkg
using Makie
using CairoMakie
using Printf
```

#### 0.1 Convolutio formulae

[ ]:
```
using Roots
using SpecialFunctions: logfactorial


# 1. BASE PROBABILITIES (Flat arrays of all 20 Amino Acids)
const FIXED_AA = Dict(
    # NNN (64 codons, 3 Stops)
    # 2 AAs with 1 codon, 9 with 2, 1 with 3, 5 with 4, 3 with 6.
    "NNN" => [fill(1/64, 2); fill(2/64, 9); fill(3/64, 1); fill(4/64, 5);␣
 ↪fill(6/64, 3)],

    # NNK/NNS (32 codons, 1 Stop)
    # 12 AAs with 1 codon, 5 with 2, 3 with 3.
    "NNK" => [fill(1/32, 12); fill(2/32, 5); fill(3/32, 3)],

    # NNB (48 codons, 1 Stop - TAG)
    # 5 AAs with 1 codon, 7 with 2, 5 with 3, 2 with 4, 1 with 5.
    "NNB" => [fill(1/48, 5); fill(2/48, 7); fill(3/48, 5); fill(4/48, 2);␣
 ↪fill(5/48, 1)],

    # DKS (12 codons, 0 Stops)
    # 8 AAs with 1 codon, 2 with 2. (Codes for exactly 10 AAs)
    "DKS" => [fill(1/12, 8); fill(2/12, 2)],

    # MAX (Ideal 20 codons, 0 Stops)
    "MAX" => fill(1/20, 20)
)

# 2. BASE RECURSION
function compositions_logdist(q::Vector{Float64}, m::Int)
    K = length(q)
```

```
    logq = log.(q)
    results = Dict{Float64,Float64}()

    function rec(i::Int, remaining::Int, sumlog::Float64, logmult::Float64)
        if i == K
            sumlog2  = sumlog + remaining * logq[K]
            logmult2 = logmult - logfactorial(remaining)
            key = round(sumlog2, digits=12)
            results[key] = get(results, key, 0.0) + exp(logmult2)
            return
        end
        @inbounds for v in 0:remaining
            rec(i+1, remaining - v, sumlog + v*logq[i], logmult -␣
↪logfactorial(v))
        end
    end

    rec(1, m, 0.0, logfactorial(m))
    return results
end

# 3. FAST CONVOLUTION & EXPONENTIATION
function convolve_logdists(d1::Dict{Float64,Float64}, d2::Dict{Float64,Float64})
    out = Dict{Float64,Float64}()
    @inbounds for (s1,m1) in d1
        for (s2,m2) in d2
            key = round(s1+s2, digits=12)
            out[key] = get(out, key, 0.0) + m1*m2
        end
    end
    return out
end

const ID_DIST = Dict{Float64,Float64}(0.0 => 1.0)

function dist_power(dist::Dict{Float64,Float64}, t::Int)
    t == 0 && return ID_DIST
    res = ID_DIST
    base = dist
    n = t
    while n > 0
        if (n & 1) == 1
            res = convolve_logdists(res, base)
        end
        n >>= 1
        n > 0 && (base = convolve_logdists(base, base))
    end
```

```julia
    return res
end

function build_dist(q::Vector{Float64}, positions::Int; block_size::Int=3)
    # Divide the total positions into manageable blocks
    b = min(block_size, positions)
    base = compositions_logdist(q, b)

    # t = number of full blocks, r = remainder
    t, r = divrem(positions, b)

    # Fast convolution multiplication
    dist = dist_power(base, t)

    # Add the remainder if it exists
    if r > 0
        remdist = compositions_logdist(q, r)
        dist = convolve_logdists(dist, remdist)
    end
    return dist
end

# 4. LIBRARY BUILDER
function build_dist_blocks(blocks::Vector{Tuple{String,Int}}; block_size::Int=3)
    first = true
    dist = Dict{Float64,Float64}()
    n_eff = 1.0
    for (mask, cnt) in blocks
        q = FIXED_AA[mask]

        # Uses the chunked builder instead of raw recursion
        d = build_dist(q, cnt, block_size=block_size)

        dist = first ? d : convolve_logdists(dist, d)
        n_eff *= length(q)^cnt
        first = false
    end
    return dist, n_eff
end

# 5. CORE PROBABILITIES (T1 and Full Coverage)
function T1_from_dist(dist::Dict{Float64,Float64}, n::Float64, S::Real)
    T1 = 0.0
    @inbounds for (sumlogp, mult) in dist
        p = exp(sumlogp)
        T1 += mult * (1 - exp(float(S) * log1p(-p)))
    end
```

```
    return T1 / n
end

function coverage_lower_bound(dist::Dict{Float64,Float64}, S::Real)
    miss = 0.0
    @inbounds for (sumlogp, mult) in dist
        p = exp(sumlogp)
        miss += mult * exp(float(S) * log1p(-p))
    end
    return max(0.0, 1.0 - miss)
end
```

```
WARNING: redefinition of constant Main.ID_DIST. This may fail, cause incorrect
answers, or produce other errors.
```

[ ]:
```
coverage_lower_bound (generic function with 1 method)
```

##### 0.1.1 Supplementary Table S1

[7]:
```
using Roots
using Printf

# 1. UNIVERSAL SOLVER (With bisection for extreme scales)
function solve_S_dist(target_prob, k, dist, n_eff)
    function get_prob(ls)
        S = 10^ls
        if k == "Full"
            return coverage_lower_bound(dist, S) - target_prob
        else
            t1 = T1_from_dist(dist, n_eff, S)
            k_num = Float64(k)
            return (1 - (1 - t1)^k_num) - target_prob
        end
    end

    # The physical brackets (1 clone to 10^20 clones)
    lower_bound, upper_bound = 0.0, 20.0

    # Boundary Checks
    if get_prob(lower_bound) >= 0
        return 1
    end

    if get_prob(upper_bound) < 0
        return -2 # Indicates S needs to be mathematically > 10^20
    end

    # Bisection is immune to flat gradients and guaranteed to converge
```

```
    try
        logS = find_zero(get_prob, (lower_bound, upper_bound), Roots.
↪Bisection())
        return round(Int, 10^logS)
    catch
        return -2
    end
end
```

[7]: solve_S_dist (generic function with 1 method)

[8]:
```
using Roots
using Printf

# 2. LaTeX FORMATTER
function latex_format(val)
    if val == -2
        return "\$>10^{20}\$"
    end

    if val < 1_000_000
        s = @sprintf("%d", val)
        return replace(s, r"(\d)(?=(\d{3})+(?!\d))" => s"\1\\,")
    else
        s = @sprintf("%.2e", Float64(val))
        parts = split(s, "e")
        base = parts[1]
        exp_val = parse(Int, parts[2])
        return "\$$base \\times 10^{$exp_val}\$"
    end
end

# 3. UNIFIED TABLE GENERATOR (L=6, 8, 10 + Mixed Scheme)

# --- TOP SECTION (L = 1, 2, 3) ---
for l in [1, 2, 3]
    for name in ["NNN", "NNB", "NNK", "MAX"]
        print("$l & $name ")
        dist, n_eff = build_dist_blocks([(name, l)])
        for p in [0.95, 0.99]
            for k in ["Full", 1, 2, 3]
                val = solve_S_dist(p, k, dist, n_eff)
                print("& $(latex_format(val)) ")
            end
        end
        println("\\\\")
    end
```

```
    println("\\midrule")
end


# --- BOTTOM SECTION (L = 6, 8, 10) ---
for l in [6, 8, 10]
    # Define the base schemes as arrays of blocks
    schemes = [
        ("NNN", [( "NNN", l )]),
        ("NNK", [( "NNK", l )]),
        ("DKS", [( "DKS", l )]),
        ("MAX", [( "MAX", l )])
    ]

    # Inject the mixed library specifically for L = 8
    if l == 8
        # We use NNK internally for NNS as they are mathematically equivalent
        push!(schemes, ("NNS:4 + DKS:4", [( "NNK", 4 ), ( "DKS", 4 )]))
    end

    for (name, blocks) in schemes
        print("$l & $name ")
        dist, n_eff = build_dist_blocks(blocks)
        for p in [0.95, 0.99]
            for k in ["Full", 1, 10, 100]
                val = solve_S_dist(p, k, dist, n_eff)
                print("& $(latex_format(val)) ")
            end
        end
        println("\\\\")
    end

    if l != 10
        println("\\midrule")
    end
end
```

```
1 & NNN & 240 & 91 & 38 & 23 & 338 & 164 & 65 & 39 \\
1 & NNB & 219 & 87 & 36 & 22 & 295 & 155 & 61 & 37 \\
1 & NNK & 173 & 79 & 36 & 23 & 223 & 129 & 58 & 37 \\
1 & MAX & 117 & 58 & 29 & 19 & 148 & 90 & 45 & 30 \\
\midrule
2 & NNN & 18\,345 & 2\,737 & 996 & 585 & 24\,627 & 5\,524 & 1\,834 & 1\,029 \\
2 & NNB & 14\,328 & 2\,461 & 900 & 533 & 18\,025 & 4\,931 & 1\,651 & 930 \\
2 & NNK & 8\,153 & 2\,129 & 875 & 533 & 9\,800 & 3\,691 & 1\,513 & 902 \\
2 & MAX & 3\,590 & 1\,197 & 598 & 399 & 4\,233 & 1\,840 & 920 & 613 \\
\midrule
3 & NNN & $1.35 \times 10^{6}$ & 79\,040 & 25\,585 & 14\,294 & $1.76 \times
```

```
10^{6}$ & 175\,835 & 50\,390 & 26\,546 \\
3 & NNB & 865\,458 & 67\,394 & 22\,069 & 12\,454 & $1.04 \times 10^{6}$ &
148\,245 & 43\,128 & 22\,886 \\
3 & NNK & 342\,436 & 55\,484 & 21\,050 & 12\,373 & 395\,173 & 102\,478 & 38\,116
& 21\,758 \\
3 & MAX & 95\,857 & 23\,964 & 11\,982 & 7\,988 & 108\,732 & 36\,839 & 18\,420 &
12\,280 \\
\midrule
6 & NNN & $4.93 \times 10^{11}$ & $1.74 \times 10^{9}$ & $3.63 \times 10^{7}$ &
$2.67 \times 10^{6}$ & $6.03 \times 10^{11}$ & $4.88 \times 10^{9}$ & $6.47
\times 10^{7}$ & $4.20 \times 10^{6}$ \\
6 & NNK & $1.92 \times 10^{10}$ & $9.32 \times 10^{8}$ & $3.06 \times 10^{7}$ &
$2.40 \times 10^{6}$ & $2.10 \times 10^{10}$ & $2.03 \times 10^{9}$ & $5.26
\times 10^{7}$ & $3.76 \times 10^{6}$ \\
6 & DKS & $4.62 \times 10^{7}$ & $5.58 \times 10^{6}$ & 335\,419 & 30\,348 &
$5.10 \times 10^{7}$ & $9.91 \times 10^{6}$ & 542\,219 & 46\,968 \\
6 & MAX & $1.34 \times 10^{9}$ & $1.92 \times 10^{8}$ & $1.92 \times 10^{7}$ &
$1.92 \times 10^{6}$ & $1.45 \times 10^{9}$ & $2.95 \times 10^{8}$ & $2.95
\times 10^{7}$ & $2.95 \times 10^{6}$ \\
\midrule
8 & NNN & $2.41 \times 10^{15}$ & $1.31 \times 10^{12}$ & $1.87 \times 10^{10}$
& $1.21 \times 10^{9}$ & $2.86 \times 10^{15}$ & $4.16 \times 10^{12}$ & $3.48
\times 10^{10}$ & $1.93 \times 10^{9}$ \\
8 & NNK & $2.52 \times 10^{13}$ & $5.94 \times 10^{11}$ & $1.47 \times 10^{10}$
& $1.05 \times 10^{9}$ & $2.69 \times 10^{13}$ & $1.42 \times 10^{12}$ & $2.61
\times 10^{10}$ & $1.65 \times 10^{9}$ \\
8 & DKS & $8.44 \times 10^{9}$ & $6.75 \times 10^{8}$ & $3.51 \times 10^{7}$ &
$3.05 \times 10^{6}$ & $9.13 \times 10^{9}$ & $1.26 \times 10^{9}$ & $5.76
\times 10^{7}$ & $4.74 \times 10^{6}$ \\
8 & MAX & $6.90 \times 10^{11}$ & $7.67 \times 10^{10}$ & $7.67 \times 10^{9}$ &
$7.67 \times 10^{8}$ & $7.31 \times 10^{11}$ & $1.18 \times 10^{11}$ & $1.18
\times 10^{10}$ & $1.18 \times 10^{9}$ \\
8 & NNS:4 + DKS:4 & $4.62 \times 10^{11}$ & $2.04 \times 10^{10}$ & $7.12 \times
10^{8}$ & $5.64 \times 10^{7}$ & $4.97 \times 10^{11}$ & $4.34 \times 10^{10}$ &
$1.22 \times 10^{9}$ & $8.82 \times 10^{7}$ \\
\midrule
10 & NNN & $>10^{20}$ & $9.58 \times 10^{14}$ & $9.74 \times 10^{12}$ & $5.55
\times 10^{11}$ & $>10^{20}$ & $3.41 \times 10^{15}$ & $1.89 \times 10^{13}$ &
$8.96 \times 10^{11}$ \\
10 & NNK & $3.14 \times 10^{16}$ & $3.73 \times 10^{14}$ & $7.07 \times 10^{12}$
& $4.58 \times 10^{11}$ & $3.32 \times 10^{16}$ & $9.74 \times 10^{14}$ & $1.30
\times 10^{13}$ & $7.32 \times 10^{11}$ \\
10 & DKS & $1.47 \times 10^{12}$ & $8.12 \times 10^{10}$ & $3.67 \times 10^{9}$
& $3.08 \times 10^{8}$ & $1.57 \times 10^{12}$ & $1.58 \times 10^{11}$ & $6.13
\times 10^{9}$ & $4.79 \times 10^{8}$ \\
10 & MAX & $3.37 \times 10^{14}$ & $3.07 \times 10^{13}$ & $3.07 \times 10^{12}$
& $3.07 \times 10^{11}$ & $3.54 \times 10^{14}$ & $4.72 \times 10^{13}$ & $4.72
\times 10^{12}$ & $4.72 \times 10^{11}$ \\
```

##### 0.1.2 Supplementary Figure S2

[5]:
```
# Calculate Tk from T1
Tk_calc(T1_val, k) = 1.0 - (1.0 - T1_val)^k

# Function to find Maximum discrete L for a given S
function find_max_L_discrete(S, target_Tk, k, schema_name; L_bounds=1:15)
    target_T1 = 1.0 - (1.0 - target_Tk)^(1/k)
    max_l = 0
    for l in L_bounds
        dist, n_eff = build_dist_blocks([(schema_name, l)])
        t1 = T1_from_dist(dist, n_eff, S)
        if t1 >= target_T1
            max_l = l
        else
            break # Once we fail the threshold, we stop checking larger L
        end
    end
    return max_l
end

# Pre-compute grids for plotting
S_test_ext = 10.0 .^ (1:0.5:15)
L_grid = 1:12
A_vals = Dict("NNN"=>20, "NNK"=>20, "DKS"=>10, "MAX"=>20) # Total distinct AAs␣
 ↪possible

# Compute T1 arrays for Panel A (Tk vs S/A^L)
T1_dict = Dict{String, Matrix{Float64}}()
for name in ["NNN", "NNK", "DKS", "MAX"]
    mat = zeros(length(S_test_ext), length(L_grid))
    for (j, l) in enumerate(L_grid)
        dist, n_eff = build_dist_blocks([(name, l)])
        for (i, s) in enumerate(S_test_ext)
            mat[i, j] = T1_from_dist(dist, n_eff, s)
        end
    end
    T1_dict[name] = mat
end
```

[ ]:
```
# FIGURA S2
fig_S2 = Figure(size=(1200, 900))

k_vals = [1, 10, 100]
styli_k = [:solid, :dash, :dot]
thr_base = 0.99
```

```
schemas = [
    ("NNN", :blue),
    ("NNK", :orange), # labeled NNS in plots
    ("DKS", :purple),
    ("MAX", :black)
]

# PANEL A: T_k vs S / A^L
axA = Axis(fig_S2[1, 1:2],
    title="A) T_k vs S / A^L", xlabel="S / A^L", ylabel="T_k",
    xscale=log10, limits=(1e-4, 1e3, -0.05, 1.05))

for (i, k) in enumerate(k_vals)
    for (idx, l) in enumerate([6, 8, 10]) # Sample specific Ls for clarity
        for (name, color) in schemas
            plot_name = name == "NNK" ? "NNS" : name
            lbl = (idx == 1 && i == 1) ? plot_name : nothing

            # Normalize sequence space
            normalized_S = S_test_ext ./ (A_vals[name]^l)
            tk_vals = Tk_calc.(T1_dict[name][:, l], k)

            lines!(axA, normalized_S, tk_vals, color=(color, 0.6),␣
 ↪linestyle=styli_k[i], label=lbl)
        end
    end
end
for (i, k) in enumerate(k_vals) lines!(axA, [NaN], [NaN], color=:black,␣
 ↪linestyle=styli_k[i], label="k = $k") end
axislegend(axA, position=:rb, nbanks=2)

# PANEL B: Max L vs S for NNS (k=10)
axB = Axis(fig_S2[2, 1],
    title="B) Max L vs S for NNS (k=10)", xlabel="S (Log10)", ylabel="Max L",
    xscale=log10, yticks=1:1:15, limits=(1e4, 1e15, 3, 14))

thresholds_T10 = [0.99, 0.95, 0.80, 0.50]
colors_thr = Makie.cgrad([:deepskyblue, :mediumblue, :midnightblue],␣
 ↪length(thresholds_T10), categorical=true, rev=true)

for (i, thr) in enumerate(thresholds_T10)
    L_nns_vals = [find_max_L_discrete(s, thr, 10, "NNK", L_bounds=1:15) for s␣
 ↪in S_test_ext]
    stairs!(axB, S_test_ext, L_nns_vals, label="T10 = $thr",␣
 ↪color=colors_thr[i], linewidth=2.5)
end
axislegend(axB, position=:lt)
```

```
# PANEL C: Max L vs S across all schemas (T=0.99)
axC = Axis(fig_S2[2, 2],
    title="C) Max L vs S (T_k = 0.99)", xlabel="S (Log10)", ylabel="Max L",
    xscale=log10, yticks=1:1:15, limits=(1e2, 1e15, 3, 14))

for (i_k, k) in enumerate(k_vals)
    for (i_s, (name, color)) in enumerate(schemas)
        plot_name = name == "NNK" ? "NNS" : name

        L_vals = [find_max_L_discrete(s, thr_base, k, name, L_bounds=1:15) for␣
 ↪s in S_test_ext]

        # Slight vertical offsets so the lines don't perfectly overlap
        offset = (i_s - 2.5) * 0.12 + (i_k - 2) * 0.04
        stairs!(axC, S_test_ext, L_vals .+ offset, color=(color, 0.6),␣
 ↪linestyle=styli_k[i_k], linewidth=2.5)
    end
end

for (name, color) in schemas lines!(axC, [NaN], [NaN], color=color, linestyle=:
 ↪solid, linewidth=3, label=(name == "NNK" ? "NNS" : name)) end
for (i_k, k) in enumerate(k_vals) lines!(axC, [NaN], [NaN], color=:black,␣
 ↪linestyle=styli_k[i_k], linewidth=2.5, label="k=$k") end
axislegend(axC, position=:lt, nbanks=2, framevisible=true)

fig_S2
# save("Figura_S2.png", fig_S2, pt_per_unit=1)
```

[ ]:

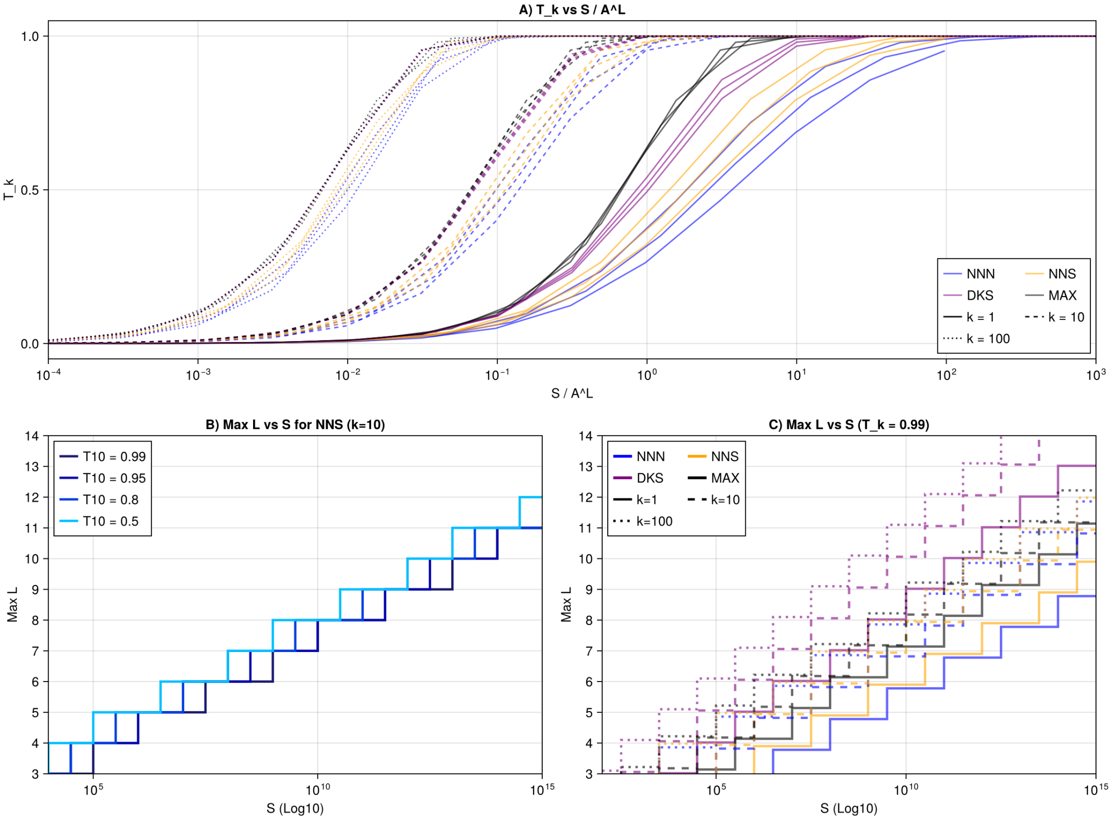
A) T_k vs S / A^L
T_k
S / A^L
NNN
NNS
DKS
MAX
k = 1
k = 10
k = 100
B) Max L vs S for NNS (k=10)
T10 = 0.99
T10 = 0.95
T10 = 0.8
T10 = 0.5
Max L
S (Log10)
C) Max L vs S (T_k = 0.99)
NNN
NNS
DKS
MAX
k=1
k=10
k=100
Max L
S (Log10)
